# Clustered VEGF Nanoparticles in Microporous Annealed Particle (MAP) Hydrogel Accelerates Functional Recovery and Brain Tissue Repair after Stroke

**DOI:** 10.1101/2025.01.30.635733

**Authors:** Kevin Erning, Katrina L. Wilson, Cara S. Smith, Long Nguyen, Neica I. Joesph, Rachel Irengo, Lauren Y. Cao, Mohanapriya Cumaran, Yi Shi, Sihan Lyu, Lindsay Riley, Timothy W. Dunn, S. Thomas Carmichael, Tatiana Segura

**Author notes:** Corresponding author: Prof. T. Segura, Department of Biomedical Engineering, Duke University, 101 Science Drive Campus Box 90281, Durham NC 27708-0281, United States.

## Abstract

Ischemic stroke, a blockage in the vasculature of the brain that results in insufficient blood flow, is one of the world’s leading causes of disability. The cascade of inflammation and cell death that occurs immediately following stroke drives vascular and functional loss that does not fully recover over time, and no FDA-approved therapies exist that stimulate regeneration post-stroke. We have previously developed a hydrogel scaffold that delivered heparin nanoparticles with and without VEGF bound to their surface to promote angiogenesis and reduce inflammation, respectively. However, the inclusion of the naked heparin nanoparticles warranted concern over the development of bleeding complications. Here, we explore how microporous annealed particle (MAP) scaffolds functionalized with VEGF coated heparin nanoparticles can both reduce inflammation and promote angiogenesis - without the inclusion of free heparin nanoparticles. We show that our updated design not only successfully promotes de novo tissue formation, including the development of mature vessels and neurite sprouting, but it also leads to functional improvement in a photothrombotic stroke model. In addition, we find increased astrocyte infiltration into the infarct site correlated with mature vessel formation. This work demonstrates how our biomaterial design can enhance endogenous regeneration post-stroke while eliminating the need for excess heparin.

## 1. Introduction

Stroke is a leading cause of serious long-term disability affecting nearly 800,000 people annually in the United States^[1]^. The most common stroke type is ischemic stroke, during which a vascular occlusion blocks the blood supply to brain tissue causing inflammation, cell death, and, in some cases, functional loss.^[2]^ Stroke mortality rates have decreased in recent years, which has been attributed to the clinical use of FDA-approved therapies, including thrombolytics, that focus on restoring blood flow and reducing the risks for additional damage.^[3]^ However, aside from physical therapy, there are no current therapeutic interventions that address the existing brain damage, life altering disabilities^[4]^, and long term secondary indications that result from stroke-brain remodeling of the increasing population of stroke survivors living with long-term disability.^[5]^

The development of therapies that increase angiogenesis, the sprouting of blood vessels from pre-existing vessels, is considered a promising strategy for enhancing regeneration and recovery post-stroke.^[6]^ Angiogenesis following stroke has significant correlation with improved functional outcomes in both animal models and human stroke patients.^[7]^ Not only does angiogenesis help reintroduce critical nutrients and oxygen to damaged tissue, but it also has been attributed to providing a permissive environment for neurogenesis, which in turn supports functional recovery – though the exact mechanisms of this are still under investigation.^[8]^ Interestingly, endogenous angiogenesis does occur following stroke, but it is limited and is associated with an increase in cells and factors associated with the inflammatory response.^[9]^ Clinical trials have already begun exploring how to leverage this endogenous post-stroke angiogenesis even further. These methods include the delivery of erythropoietin, an angiogenic hormone, in combination with a multiple burr hole surgery through the skull and dura mater to increase revascularization (NCT02603406)^[10]^, as well as the delivery of human urinary kallidinogenase (NCT03431909), a proteinase extracted from urine that has been previously shown to promote angiogenesis in an middle cerebral artery occlusion (MCAO) rat model.^[11]^ Some clinical trials are also evaluating how to better evaluate angiogenesis in the brain in preparation to assess new therapies targeting this mechanism for stroke treatment (NCT01656785).^[12]^ However, while promising, these current strategies have several disadvantages, including non-localized delivery of therapeutic agents and a reliance on using inflammatory procedures to induce revascularization from vessels external to the brain tissue. Furthermore, many of these strategies, particularly those that rely on tissue disruption to induce revascularization, do not consider how the introduction of additional inflammation may hinder other aspects of the regenerative process post-stroke. Thus, future angiogenic therapies must balance how to both stimulate blood vessel formation while still considering other facets of the injury response, such as inflammation.

Vascular endothelial growth factor (VEGF), which plays important roles in the development and maintenance of both the circulatory and nervous systems^[13]^, is also a major factor in the injury response following ischemic stroke, with complex effects on several mechanisms, including angiogenesis, neurogenesis, inflammation, and blood-brain barrier maintenance.^[14]^ Following ischemic stroke, VEGF and its receptors have been shown to be upregulated near and within the infarct site^[15]^, which is thought to contribute to the growth of new vessels near the border of the infarct site within 72 hours after injury^[15b]^. The delivery of exogenous VEGF has similar effects. When infused over the course of 1 week into the cortex of healthy rats, soluble VEGF was shown to significantly increase angiogenesis - however, it also increased the proliferation of activated astrocytes and blood-brain barrier permeability.^[16]^ In the context of stroke, VEGF has been delivered intranasally^[17]^ and intracerebroventricularly^[18]^, resulting in functional recovery in rodent stroke models. Another recent study considered whether a microneedle-based delivery systems could improve VEGF delivery through better localization of the therapeutic, and found significant improvement in both angiogenesis and neurogenesis in *in vivo* rodent models.^[19]^ However, while these VEGF delivery strategies have shown promise, they rely upon the delivery of multiple or continuous doses of soluble VEGF, increasing both the therapeutic cost and the risk of potential off target effects.

Hyaluronic acid (HA)-based hydrogels, which have already been shown to enhance wound healing^[20]^, cartilage regeneration^[21]^, and brain regeneration^[22]^, are a promising platform to deliver better targeted and potent angiogenic therapies. These hydrogels are easily injected into the injury site and can gel *in situ*, making them localized and minimally invasive, as well as able to completely fill an injury site.^[23]^ Already, HA-based hydrogels have been used to deliver bioactive cues such as stem cells, growth factors, or small molecule drugs into the infarct site post-stroke to promote angiogenesis.^[24]^ In an effort to further improve upon these HA-based scaffold systems, we have previously pioneered HA-based microporous annealed particle (MAP) scaffolds, which are comprised of crosslinked individual HA microgels that allow for increased cellular infiltration and migration through the void spaces between microgels.^[25]^ They have also been successfully utilized in platforms targeting brain regeneration, including as a stroke treatment.^[26]^

Here, we explore using HA-based MAP as a delivery vehicle for VEGF. Our design only utilizes VEGF coated heparin nanoparticles, improving upon a previous design in which both VEGF coated and naked heparin nanoparticles were co-delivered within a traditional bulk HA hydrogel to target angiogenesis and inflammation, respectively.^[27]^ While this previous strategy was effective in mediating inflammation and regeneration post-stroke, it raised concerns over the use of an exposed heparin surface, which could potentiate anticoagulant effects. We hypothesize that the use of MAP, which have a known anti-inflammatory effect following stroke, as our VEGF coated heparin nanoparticle vehicle will reduce inflammation and promote angiogenesis, neurogenesis, and functional recovery, eliminating the need for naked heparin nanoparticles. To demonstrate this, we first confirm that our updated heparin nanoparticle design does not have an anticoagulant effect *in vivo* at therapeutically relevant concentrations. Then, *in vitro*, we confirm the ability of our VEGF heparin nanoparticle design to activate VEGF receptors. We then compare MAP containing VEGF coated heparin nanoparticles alone and MAP containing both VEGF heparin nanoparticles and naked heparin nanoparticles in a mouse photothrombotic stroke model. We ultimately demonstrate that MAP containing VEGF coated heparin nanoparticles alone had the most significant impact on mature vessel formation and neurite sprouting within the infarct and functional recovery.

## 2. Results and Discussion

### 2.1. VEGF modified Heparin-Norbornene Nanoparticles result in Long-lasting MAP Scaffold Tethering and enhanced VEGF receptor-2 phosphorylation

Heparin, a highly sulfated glycosaminoglycan, has the capacity to bind various different growth factors, including VEGF.^[28]^ As such, it has been used extensively in biomaterial design to bind, protect, and ultimately deliver growth factors that would otherwise quickly disperse and degrade *in situ*.^[29]^. However, because unpolymerized heparin is a known anti-coagulant, we chose to utilize heparin nanoparticles (nH) in our biomaterial design. nH has been previously shown to not have the same anti-coagulatory effects as unpolymerized heparin.^[27]^ In addition, the use of nH as a protein-binding substrate helps to both cluster multiple proteins together and alter the presentation of the bound protein to cell receptors, both of which have been previously demonstrated to improve bioactivity.^[26b, 27, 30]^ Inverse emulsion polymerization of hexane and water was used to synthesize our heparin nanoparticles (nH) using two different chemistries: (1) radical polymerization with heparin-methacrylamide-ABH (ρ-Azidobenzoyl Hydrazide) as the reactive component and APS/TMED as the radical initiator (nH-ABH) and (2) UV initiated thiol-ene click chemistry with heparin-norbornene/dithiothreitol (DTT) as the reactive components and LAP as the reaction initiator, as previously described (nH-NB) (**Figure 1A**).^[26b, 27]^ nH-NB was measured 93 nm in diameter as by scanning electron microscopy (SEM) (**Figure 1B**). With dynamic light scattering (DLS), both nH-ABH and nH-NB were approximately 100 nm in diameter (**Figure 1C**). To immobilize VEGF to the surface of the nH-ABH, VEGF was reacted with the ABH groups on the heparin backbone via UV irradiation to form a covalent bond.^[30]^ For nH-NB, LAP activation of a nH-NB/VEGF solution resulted in robust and stable covalent conjugation of VEGF to the nanoparticle surface. VEGF binding efficiency (at a ratio of 20 ng VEGF:1 ng nH) for nH-ABH and nH-NB was measured by evaluating the amount of VEGF washed away from the nanoparticles using a VEGF ELISA (R&D Systems) (**Figure 1D**), and we found that nH-NB allowed for consistent 99.99% binding efficiency, compared to an average of 76.84% efficiency for nH-ABH (**Figure 1E**). Finally, to verify that the new thiol-ene chemistry does not alter VEGF bioactivity, we performed a VEGF receptor 2 (VEGFR-2) phosphorylation assay on VEGF coated nH-ABH and nH-NB treated human umbilical vein endothelial cells (HUVECs). 15 mins after treatment, we found that VEGF bound to nH-ABH and nH-NB formulations activated VEGFR-2 as well as and better, respectively, than unmodified soluble VEGF (**Figure 1F**). Interestingly, clustered VEGF on nH-NB yielded a more elevated phosphorylation than soluble VEGF and clustered VEGF on nH-ABH despite the same amount of VEGF being used. We suspected this is due to a higher VEGF clustering density in nH-NB compared to nH-ABH as we have shown that higher clustering resulted in more *de novo* vessels *in vivo*.^[31]^

**Figure 1.**
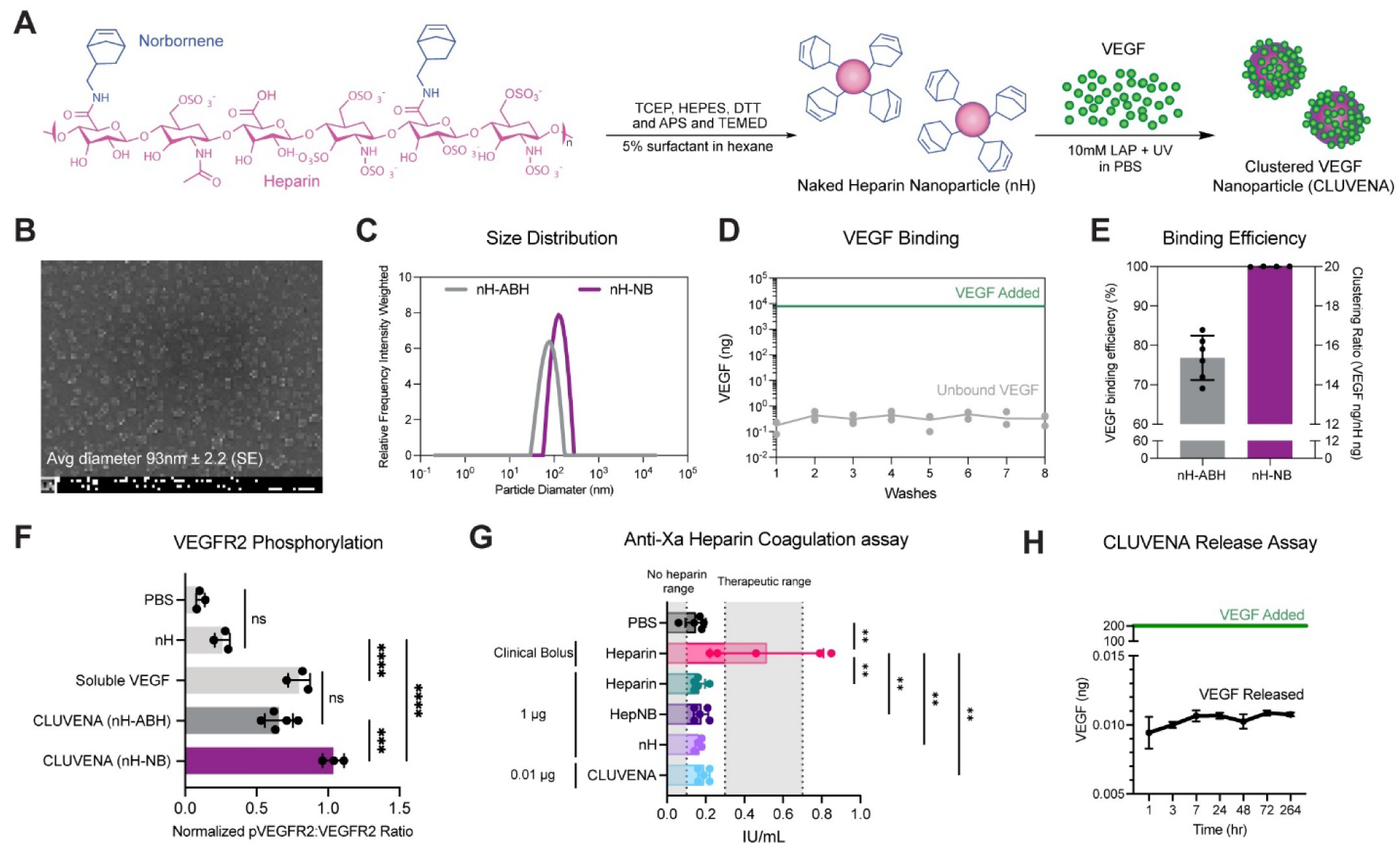
Fabrication and characterization of heparin nanoparticles (nH) covalently functionalized with VEGF. A) Schematic of **CLU**stered **VE**GF **NA**noparticles (CLUVENA) production. Norbornene-modified heparin was processed into heparin-norbornene nanoparticles (nH) after which VEGF was covalently bound to their surface. B) Representative scanning electron microscopy (SEM) image of gold coated nH-NB. C) Dynamic Light Scattering (DLS) data comparing the relative particle size distribution between nH-ABH and nH-NB nanoparticles. D) VEGF ELISA results detailing the amount of unbound VEGF collected in wash solutions following the covalent binding of VEGF with nH-NB. E) VEGF binding efficiency between heparin-ABH and heparin-NB nanoparticles as quantified by VEGF ELISA of unbound VEGF in flowthrough and washes. F) Quantification of the normalized pVEGFR2/VEGFR2 ratio in HUVECs by ELISA. HUVECs were treated for 15 min. G) Quantification of VEGF ELISA to determine the amount of VEGF released from a CLUVENA-MAP scaffold over 264 hrs. H) Anti-Xa heparin coagulation assay. For panels F and H, one-way ANOVA with a Tukey HSD post-hoc analysis was performed. * *p* < 0.05, ** *p* < 0.01, *** *p* < 0.001, **** *p* < 0.0001. Error bars represent standard deviation.

While the ability of heparin to bind growth factors such as VEGF is highly valuable in biomaterial design, heparin is also widely used clinically as an anti-coagulant, raising concerns of potential off-target effects. Heparin primarily interferes with the coagulation cascade by inactivating thrombin (factor IIa) and activated factor X (factor Xa) via binding to antithrombin (AT) and its inhibitory activity.^[32]^ Heparin binds to AT with its pentasaccharide sequence, leading to AT undergoing a conformational change that makes it orders of magnitude more reactive to thrombin and factor Xa.^[33]^ Because previous nanoparticle formulations of heparin have been shown to negate the ability of heparin to impair blood clotting^[27]^, we hypothesized that our updated HA-NB and CLUVENA formulations would behave similarly *in vivo*. To confirm this, we ran a standard clinical heparin coagulation test, an anti-Xa assay, on mice injected with heparin-norbornene and CLUVENA (**Figure 1G**). An anti-Xa assay, also known as a heparin assay, indirectly measured how much of our injected treatment behaves as unmodified porcine intestinal unfractionated heparin (UFH) by measuring its inhibition of clotting factor Xa activity. Promisingly, the covalent modification of VEGF onto the surface of HA-NB (CLUVENA) did not influence blood clotting, with significantly lower Factor Xa inhibition in comparison to heparin and similar values to that of our 1x PBS negative control. Therefore, because of its reduced risk of blood thinning, we next sought to confirm whether the addition of CLUVENA alone into MAP can influence recovery post-stroke.

Due to the superior performance of VEGF coated HA-NB *in vitro* (which will now be referred to as **CLU**stered **VE**GF **NA**noparticles (CLUVENA)), we moved forward with fabricating hyaluronic acid (HA)-based microporous annealed particle (MAP) scaffolds with CLUVENA covalently attached to its surface for further study. HA hydrogel microparticles (microgels) 100 μm in diameter were first fabricated through a microfluidic flow focusing device crosslinking HA-NB polymers with an MMP degradable dithiol peptide and LAP/UV light as the radical initiators (**Figure S1A-C**). Per our previous study, microgels were also decorated with RGD (1 mM) to promote astrocyte infiltration.^[26a]^ To form an interconnected MAP scaffold network, 4arm-PEG-tetrazine was mixed with the microgels and allowed to anneal. The storage modulus of the resulting HA-NB MAP scaffolds have been previously established to range from 150 – 5,800 Pa, depending on the ratio of PEG-tetrazine crosslinker to HA.^[34]^ In these studies, we used the previously optimized Tet/HA ratio of 7, which resulted in a storage modulus of approximately 300 Pa (**Figure S1D-E**), because of its similarity to the brain’s storage modulus.^[34–35]^ Furthermore, we established that our MAP scaffolds have a void volume fraction of about 25% (**Figure S1F-G**). MAP scaffold 3D pores were also analyzed with LOVAMAP (**Figure S1H-L**).^[36]^ These results showed that our MAP scaffolds had pores that could accommodate rigid sphere diameters ranging from 10 – 50 μm (**Figure S1I**), with a median 3D pore volume of 34 pL (**Figure S1J-K**) and a median 3D pore aspect ratio of 3.83 (**Figure S1L**). CLUVENA was loaded into MAP during the microgel annealing process at 200 ng VEGF per 6 μL HA microgels, which resulted in MAP containing covalently attached CLUVENA. A release assay in PBS at 37 °C showed no release of VEGF (quantified with ELISA) throughout the duration of the study (**Figure 1H**), confirming successful CLUVENA loading onto the MAP scaffold.

### 2.2. MAP Scaffold Attenuates Astrocyte Reactivity Despite VEGF Delivery

In our original report to treat ischemic stroke, an HA hydrogel containing both CLUVENA and nH, to target angiogenesis and inflammation, respectively, was found to reduce astrogliosis, induce neurite sprouting, and enhance functional recovery.^[27]^As expected because of its anti-inflammatory effects^[37]^, removing nH from this design resulted in a loss of astrogliosis attenuation, no neurite sprouting, and elevated inflammation.^[27]^ Thus, the inclusion of naked nH was deemed essential for this therapeutic intervention. However, given the risks associated with heparin, we sought to design an HA-based hydrogel system that could still induce angiogenesis, neurogenesis, and functional recovery and reduced inflammation - without the inclusion of excess naked nH. Because we have previously found that MAP delivered into the stroke core resulted in the attenuation of post-stroke astrogliosis^[38]^ and astrocyte and microglia/macrophage reactivity^[26a]^, we reasoned that the delivery of CLUVENA alone in MAP could similarly simultaneously target angiogenesis and inflammation, without the need for additional nH. To validate this hypothesis, we sought to evaluate MAP loaded with either CLUVENA alone or CLUVENA + nH, in an *in vivo* stroke model and evaluate their ability to influence angiogenesis, neurogenesis, inflammation, and functional recovery (**Figure S2**).

Prior to histological and functional assessment of MAP treated mice, we first optimized a photothrombotic (PT) stroke model, which induces thrombi formation by photo-activation of an injected light sensitive dye that releases reactive oxygen species while mice are under anesthesia, to evaluate our treatments.^[39]^ We initially considered how isoflurane levels as well as the length of laser irradiation could impact the infarct size, measured by glial barrier formation (**Figure S3A-B**). Interestingly, we found significant differences in the infarct volume depending on the percentage of isoflurane administered during the stroke procedure (**Figure S3C-D**). The infarct volume was significantly larger at 1.5% isoflurane compared to 2.0% (**Figure S3E**). We attribute this to the fact that isoflurane is a known vasodilator, including in cerebral vasculature^[40]^, and can also effect PT stroke induction, as a study showed that PT stroke induced in a conscious mouse resulted in a significantly larger infarct compared to mouse under isoflurane anesthesia^[41]^. Moving forward with 1.5% isoflurane, we next considered the length of laser irradiation’s effect on infarct volume and determined that 13 mins resulted in the most consistent strokes (**Figure S2F-G**). With these parameters, we next sought to investigate differences in PT stroke between male and female mice (**Figure S2H-J**). Notably, we found that female mice had significant variability in stroke incidence rate and size in comparison to male mice, with ∼50% having no or small strokes and ∼25% having significantly larger strokes (**Figure S2J**). These results are supported by prior studies, in which variable stroke outcomes in young female murine experimental models are attributed to sex related differences, such as the expression of neuroprotective hormones such as estrogen.^[42]^ Therefore, we decided to use only male mice in our PT stroke studies.

For our optimized *in vivo* experiments, mice were treated with PBS, MAP, MAP+CLUVENA (M+C), or MAP+CLUVENA+nH (M+C+N) via stereotactic injection into the stroke cavity 5 days post-stroke. Mice were then sacrificed at day 15 and day 35 post-stroke for histological evaluation (**Figure 2A**). IHC staining of coronal sections revealed that the degraded gel area showed no significant differences across all three gel groups (**Figure 2B,F**). Scar thickness was quantified by measuring the shortest radial distance from infarct boundary to GFAP+ cells with elongated reactive astrocyte morphology, excluding the corpus collosum. We found that MAP injection resulted in significantly lowered scar thickness as early as day 15 compared to stroke only (**Figure 2C,G**). In addition, there was a significant increase in astrocyte residency within the MAP scaffold (**Figure 2D,H**) and astrocyte infiltration distance from infarct boundary (**Figure 2E,I**), regardless of gel treatment. Overall, MAP was able to attenuate astrogliosis and promote astrocyte infiltration despite co-delivery of proinflammatory VEGF, which we attribute to MAP’s well established anti-inflammatory effect.^[27]^

**Figure 2.**
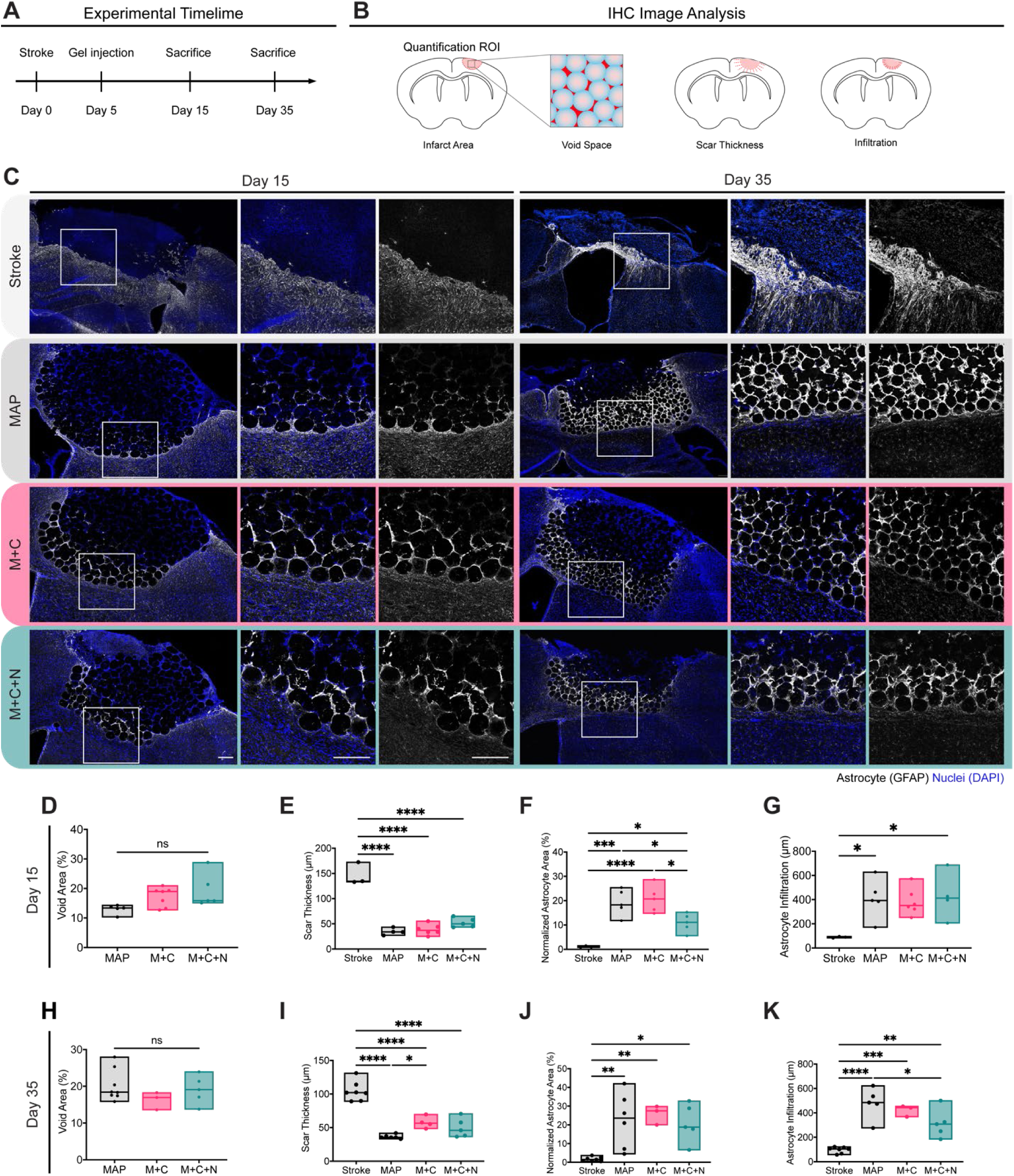
CLUVENA-MAP influences astrogliosis and astrocyte infiltration post-stroke. A) Experimental timeline and image analysis schematic. B) Representative immunofluorescent 20x images stained for astrocytes (GFAP) in white and nuclei (DAPI) in blue. Images at day 15 and 35 were quantified for C) void space, D) astrocytic scar thickness, E) normalized astrocyte area, and F) astrocytic infiltration into infarct/gel. Each data point is a biological replicate averaged from two coronal sections and plotted in a floating bar (min to max). *n* = 3 – 6. For panels B-I, one-way ANOVA with a Tukey HSD post-hoc analysis was performed. * *p* < 0.05, ** *p* < 0.01, *** *p* < 0.001, **** *p* < 0.0001. Scale bars represent 200 μm.

### 2.3. CLUVENA is Essential in Significantly Increasing *de novo* Perfused Vasculature in Infarct

We have previously established that CLUVENA alone increases angiogenesis with or without nH.^[27]^ As such, we similarly analyzed mice sacrificed at day 15 and 35 post-stroke for angiogenesis (**Figure 3A-B**). Tomato lectin (TL), injected intravenously prior to tissue processing, was used as a marker for functional and perfused blood vessels. Tomato lectin binds to the endothelial glycocalyx, a gel layer comprised of glycosaminoglycans, proteoglycans, and adsorbed plasma proteins, within the lumen of a blood vessel.^[43]^ Others have demonstrated that the glycocalyx plays an indispensable role in blood-brain barrier (BBB) maintenance.^[44]^ For example, the glycocalyx maintains barrier function in cerebral blood vessels and its degradation has led to high BBB permeability and cerebral edema.^[45]^ Therefore, we hypothesized that glycocalyx density could be used as a marker for BBB and vascular health and maturity. As early as 15 days post-stroke, we saw a significant increase in perfused vessels area in the infarct with CLUVENA delivery (M+C+ and M+C+N) compared to MAP (**Figure 3C**). CLUVENA also accelerated vessel infiltration into the center of the infarct as early as day 15 (M+C and M+C+N) compared to MAP (**Figure 3D**). Interestingly, between day 15 and 35, only M+C had increased perfused vessels in the infarct, whereas MAP and M+C+N did not (**Figure 3D**).

**Figure 3.**
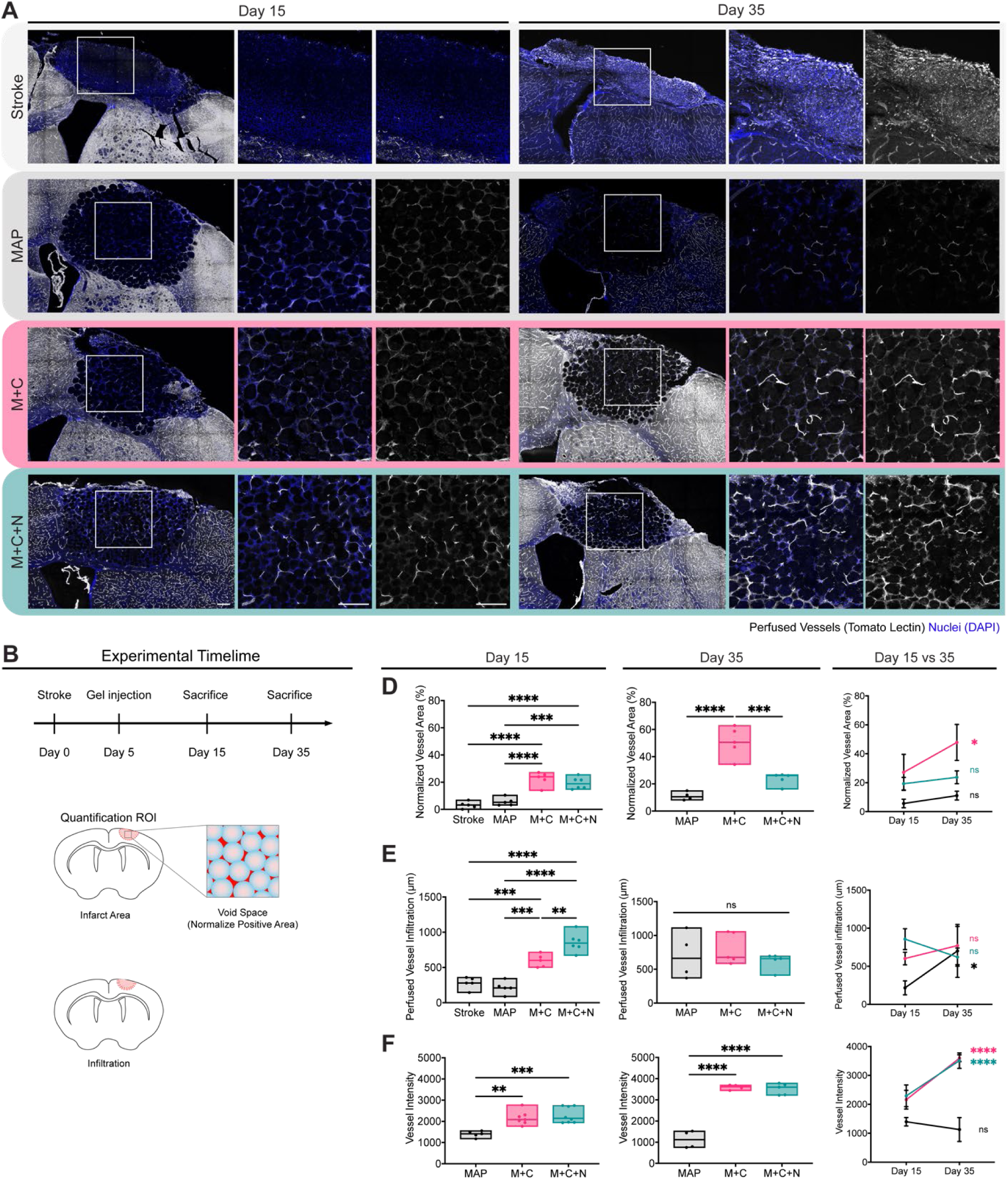
Clustered VEGF nanoparticles (CLUVENA) promote *de novo* formation of perfused vessels. A) Representative immunofluorescent 20x images stained for perfused vessels (Tomato Lectin) in white and nuclei (DAPI) in blue. B) Experimental timeline and image analysis schematic. Day 15 and 35 quantification of C) normalized perfused vessel area, D) perfused vessel infiltration, and vessel intensity as well as E) direct comparison between day 15 and 35. For panels C-E representing one timepoint, each data point is a biological replicate averaged from two coronal sections and plotted in a floating bar (min to max). For panels C-E comparing day 15 and 35, averaged values are plotted with error bars represent s.d.. *n* = 4 – 6. For panels C-E, one-way ANOVA with Tukey HSD post-hoc analysis was performed. To compare across time within treatment groups, two-way ANOVA with Sidak post-hoc analysis was performed. * *p* < 0.05, ** *p* < 0.01, *** *p* < 0.001, **** *p* < 0.0001. Scale bars represent 200 μm.

CLUVENA (M+C and M+C+N) significantly increased glycocalyx density in both day 15 and 35 when compared to MAP alone (**Figure 3E**). Specifically, delivery of CLUVENA resulted in further increase of glycocalyx density between day 15 and 35 compared to MAP. Overall, CLUVENA delivered with MAP promoted formation of *de novo* functional perfused vessels and BBB maturation. Moreover, MAP+CLUVENA alone resulted in the most perfused vessels at day 35.

### 2.4. MAP+CLUVENA Increases Perivascular AQP4

In the brain, *de novo* vessels require the establishment of a blood-brain barrier (BBB) for proper homeostatic function.^[46]^ Given that we observed a more abundant glycocalyx with M+C and M+C+N, we further explored BBB formation with a marker for astrocytic end-feet, another critical component of proper blood-brain barrier formation.^[47]^ We specifically assessed the expression of aquaporin-4 (AQP4), a water channel protein that links perivascular astrocytic end-feet to the BBB.^[48]^ Beyond its homeostatic function, AQP4 has also been shown to be elevated in the peri-infarct region after ischemic stroke and in areas of glial-specific swelling during cerebral edema.^[49]^ This increase in AQP4 is corroborated by our findings, as shown by our images in the peri-infarct region and boundary of the gel (**Figure 4A**). Given that AQP4 is elevated during both cerebral edema and vessel maturation, analysis of vessel maturity and BBB coverage was performed by measuring AQP4 signal that overlaps with TL signal (**Figure 4B**). Analysis showed that there were no significant differences between all experiment conditions at day 15 (**Figure 4C**). However, at day 35, both M+C and M+C+N showed a significant increase compared to MAP alone. Promisingly, only M+C showed a significant increase between day 15 and 35 (**Figure 4C**). Higher magnification images at 60x objective to highlight overlap between AQP4 and TL (**Figure 4D**).

**Figure 4.**
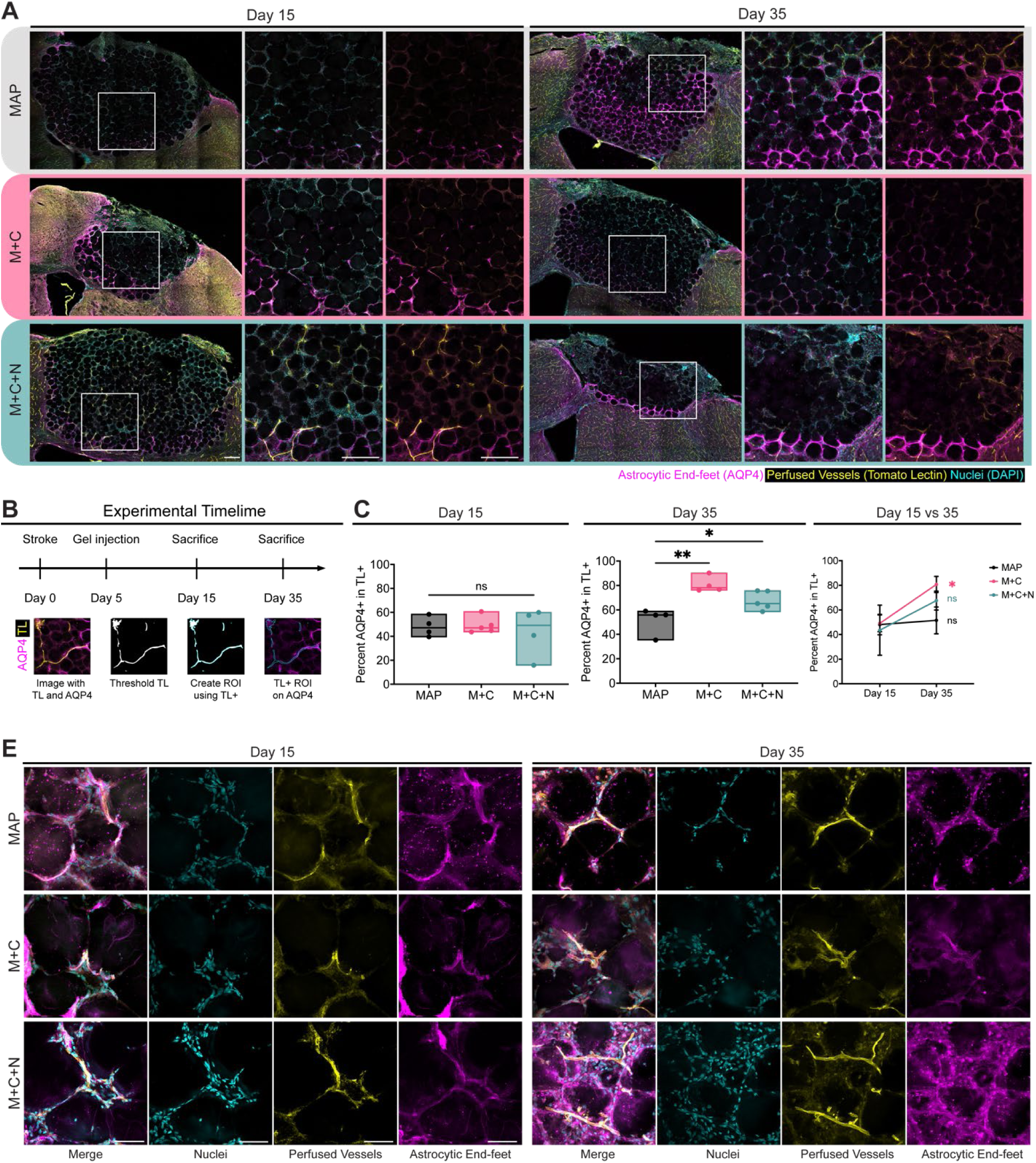
Astrocytic end-feet overlap *de novo* vessels within MAP. A) Representative immunofluorescent 20x images stained for astrocytic end-feet (AQP4) in purple, perfused vessels (Tomato Lectin) in yellow and nuclei (DAPI) in cyan. B) Experimental timeline and image analysis schematic. C) AQP4 coverage on TL in infarct on days 15 and 35, and a direct comparison of AQP4 coverage on TL between days 15 and 35. D) Immunofluorescent 60x images stained for astrocytic end-feet (AQP4) in purple, perfused vessels (Tomato Lectin) in yellow and nuclei (DAPI) in cyan. For panels C representing one timepoint, each data point is a biological replicate averaged from two coronal sections and plotted in a floating bar (min to max). For panels C comparing day 15 and 35, averaged values are plotted with error bars represent s.d. *n* = 4 – 5. For panels B-C, one-way ANOVA with a Tukey HSD post-hoc analysis was performed. To compare across time within treatment groups, two-way ANOVA with Sidak post-hoc analysis was performed. * *p* < 0.05, ** *p* < 0.01, *** *p* < 0.001, **** *p* < 0.0001. Scale bars in panel A represent 200 μm and in panel D represent 50 μm.

### 2.5. MAP-CLUVENA Alone Stimulates Neurite Elongation into the Infarct and Improves Functional Recovery

The focus of our MAP therapy development is to ultimately address the current unmet need of restoring functional loss following an ischemic event. Therefore, we also wanted to assess neurite sprouting and any corresponding functional recovery in MAP treated mice post-stroke. First, we examined neurite sprouting by considering the amount of the neuronal cytoskeletal marker neurofilament 200 (NF200) positive area within the MAP treated infarct region at day 15 and day 35. (**Figure 5A-B**). MAP treatments with CLUVENA (M+C and M+C+N) both had a significantly increased level of NF200 within the infarct when compared to MAP alone at both day 15 and day 35, suggesting that the presence of VEGF increases neurite sprouting (**Figure 5C-D**). Promisingly, neurite sprouting significantly increased in both M+C and M+C+N treated mice from day 15 to day 35 (**Figure 5E**). In contrast, mice treated with MAP alone (no CLUVENA) showed a significant decrease from day 15 to day 35, resulting in a level of neurite sprouting similar to that of the no treatment group (**Figure 5E**).

**Figure 5.**
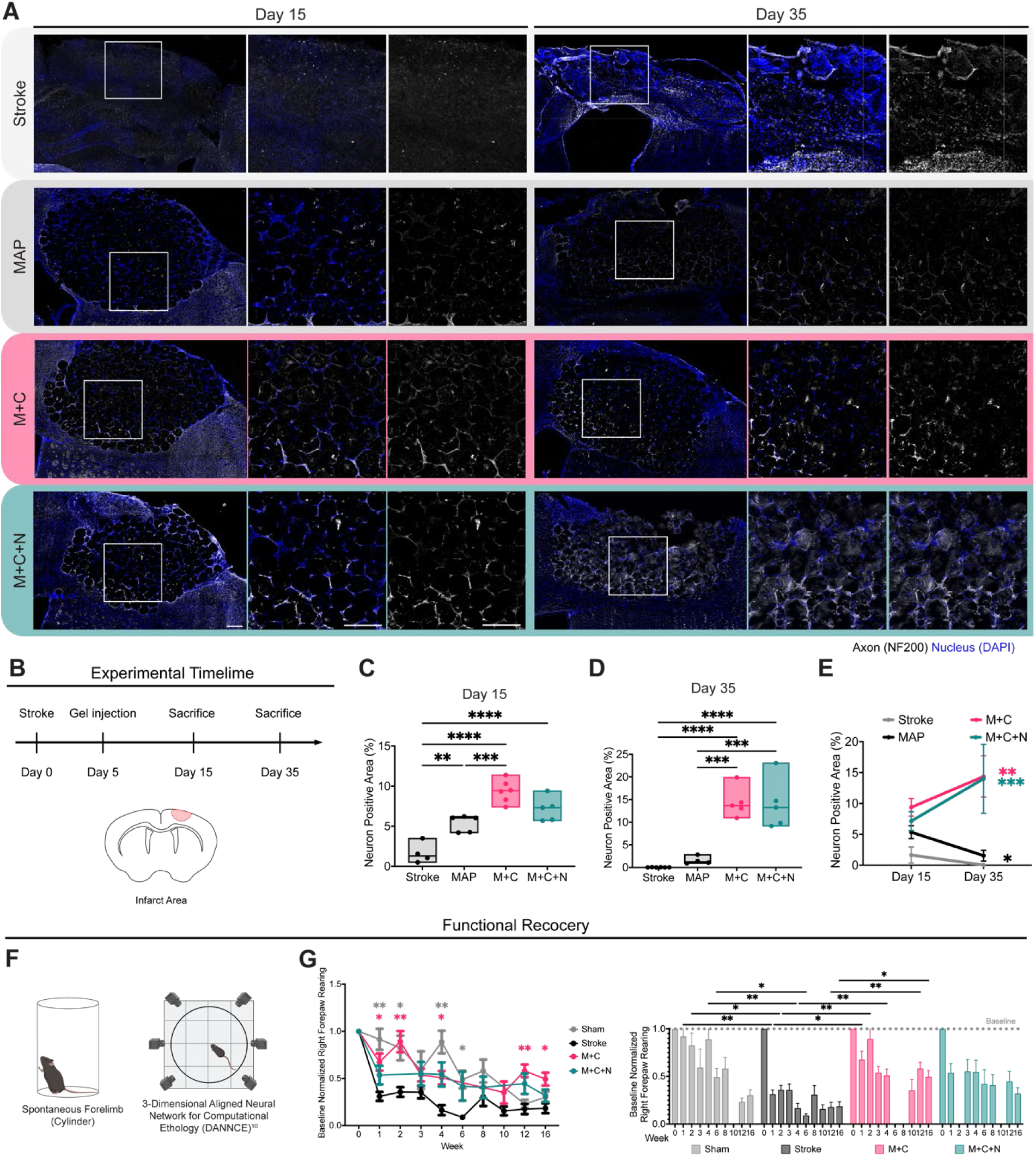
CLUVENA-MAP improves neurite sprouting and functional recovery post-stroke. A) Representative immunofluorescent 20x images stained for neurons (NF200) in white and nuclei (DAPI) in blue. B) Experimental timeline and image analysis schematic. Quantification of C) Day 15 and D) Day 35 neuron positive area as well as E) a direct comparison of neuron positive area between days 15 and 35. F) Spontaneous forelimb or cylinder test depicting contralateral right forepaw rearing activity. G) 3-Dimensional Aligned Neural Network for Computational Ethology (DANNCE) quantification of the baseline normalized right forepaw rearing represented by both a line and bar graph for sham, stroke, M+C, and M+C+N conditions over 16 weeks. For panels C-D, one-way ANOVA with a Tukey HSD post-hoc analysis was performed. For panel E, to compare across time within treatment groups, two-way ANOVA with a Sidak post-hoc analysis was performed. For panels C-D, each data point is a biological replicate averaged from two coronal sections and plotted in a floating bar (min to max). For panel E, comparing day 15 and 35, averaged values were plotted with error bars representing s.d.. *n* = 4 – 6. For panel G, mixed-effect analysis was performed with a Sidak post-hoc analysis. Error bars represent s.e.m. n = 5 – 8. * *p* < 0.05, ** *p* < 0.01, *** *p* < 0.001, **** *p* < 0.0001. Scale bars represent 200 μm.

Given the significant improvement in neurite sprouting in CLUVENA containing treatment groups in comparison to both MAP alone and no treatment groups, we hypothesized that CLUVENA containing MAP treatments would correlate with an increase in functional recovery. Because our PT stroke model induces stroke in the left motor cortex, we specifically measured behavioral changes in the contralateral right forelimb. We performed the spontaneous forelimb or cylinder test, assessing right forepaw deficit and functional recovery (**Figure 5F**). We employed a 3-dimensional animal pose tracking technology called 3-Dimensional Aligned Neural Network for Computational Ethology (DANNCE) to precisely track and measure the time the contralateral forepaw rested on the glass cylinder during a rearing event.^[50]^ Because CLUVENA containing MAP improved not only neurite sprouting but also astrogliosis, angiogenesis, and vessel maturation, we expected both M+C and M+C+N to exhibit comparable degrees of functional recovery. Surprisingly, however, only M+C exhibited any significant functional recovery (**Figure 5F**). Moreover, M+C treated mice showed significant functional improvements in comparison to the stroke control mice as early as week 1 (2 days post gel injection) (**Figure 5F**). While there were no significant differences in neurite sprouting to explain the functional differences between M+C and M+C+N, M+C did have a significantly higher normalized vessel area within the infarct site (**Figure 3D**), suggesting that the functional behavior could be correlated to vascularization. Another hypothesis we have is that the heparin nanoparticles, which have been previously demonstrated to improve functional recovery in a stroke model when delivered in a solid HA hydrogel^[27]^, are more exposed to diffusing proteins and other factors within the highly porous MAP scaffold that could bind to their surface, altering their bioactive effect. Many prior heparin nanoparticle and heparinized scaffold designs have their heparin element pre-loaded with a specific peptide or protein of interest because of the negatively charged heparin structure’s high binding affinity for an array of different molecules in the body.^[29a, 51]^ Thus, perhaps rather than binding proteins that could support functional recovery, our naked heparin nanoparticles may be binding proteins that neutralize or alter some of its therapeutic effect.^[52]^

## 3. Conclusion

Here, we discuss the development, characterization, and implementation of an improved **CLU**stered **VE**GF **NA**noparticle (CLUVENA) design that is ultimately integrated into a hyaluronic acid-based microporous annealed particle (MAP) scaffold to improve tissue regeneration and functional recovery post-stroke. The fabrication of CLUVENA used in these studies utilized a norbornene-functionalized heparin to covalently bind VEGF to its surface, improving both its overall VEGF binding efficacy and its VEGF receptor 2 activation *in vitro*. Interestingly, with an anti-coagulation assay, we found that CLUVENA had no observable anti-coagulatory effect, with similar values of Factor Xa present in the blood as our negative control at therapeutically relevant concentrations. When CLUVENA was incorporated into MAP scaffolds, VEGF was retained, which we considered promising for further *in vivo* evaluation.

Using an optimized photothrombotic (PT) stroke mouse model, we examined whether MAP + CLUVENA scaffolds could successfully leverage both the regenerative effects of VEGF and the known properties of MAP alone^[25–26]^ to promote tissue regeneration and functional recovery. In these experiments, in addition to considering MAP alone and no treatment groups, we also compared MAP + CLUVENA (M+C) to MAP + CLUVENA + naked heparin nanoparticles (nH) (M+C+N). Histological assessment of the infarct site 15 and 35 days after stroke revealed that all CLUVENA containing MAP treatments reduced glial scarring and increased astrocyte and vessel infiltration as well as neurite sprouting. Surprisingly, however, we found that only M+C showed significant functional recovery, despite lacking the additional therapeutic component of naked heparin nanoparticles. From our histological assessment, the only significant difference between M+C and M+C+N treatments was the quantity of perfused vessels within the infarct site – M+C had significantly more than M+C+N at the later timepoint. While this suggests that the functional differences could be related to vessel perfusion within the infarct site, future studies should consider whether the naked heparin nanoparticle’s bioactivity is being influenced by either its inclusion within MAP or by its surrounding environment. For example, the naked heparin nanoparticles in our system may be non-specifically binding other molecules in the stroke injury environment that interfere with its effect, ultimately leading to less functional recovery. Overall, our work demonstrates the promise of utilizing CLUVENA in MAP scaffolds for not only stroke and other central nervous system related therapeutics but also for other tissue engineering applications that necessitate enhanced vascularization.

## 4. Experimental Section/Methods

### Heparin-ABH Synthesis

Heparin was modified with *p-*azidobenzoyl hydrazide and methacrylamide as previously described.^[27]^ 50 mM 2-(N-morpholino(ethansulfonic acid) (MES) buffer (pH 5.5) was used to dissolve 100 mg heparin from porcine intestinal mucosa (Alfa Aesar, Haverhill, MA, A16198) in the presence of 165 mg of 4-(4,6-dimethoxy-1,3,5-triazin-2-yl)-4-methylmorpholinium chloride (DMTMM) (TCI America, Portland, OR) and 1.5 mL of *p-*azidobenzoyl hydrazide (ABH) (BioWorld, Dublin, OH) in the dark at room temperature. After 2 hr, 537 mg of N-(3 Aminopropyl) methyacrylamide hydrochloride (APMA) (Ambeed, Arlington Heights, IL) and 365 mg of DMTMM was added to the reaction and allowed to react overnight in the dark at room temperature. Hep-ABH-MA was purified by dialysis (6 – 8 K MW) against DI water for 3-4 days and lyophilized, after which it was stored at -20°C until use. H’ NMR was taken to determine percent modification. To determine functionalization, proton NMR shifts in D_2_O of APMA at δ5.60 or δ5.75 (1H), and ABH at δ7.20 or δ7.80 ppm (2H), were compared to the heparin methyl group δ2.00 ppm. The moles of the heparin repeat unit were used to determine all equivalents.

### Heparin-NB Synthesis

Heparin was modified with norbornene (NB) as previously described.^[26b]^ Briefly, 20 mL 50 mM (MES) buffer (pH 5.5) was used to dissolve 1 g of heparin from porcine intestinal mucosa (Alfa Aesar, Haverhill, MA, A16198), whereby its carboxylic acids were activated by 1.472 g of DMTMM and allowed to react for 10 min at room temperature with stir bar. 147 μL norbornene-2-methylamine (NMA) (TCI America, Portland, OR, N0907) was slowly added while stirring vigorously. The reaction was then capped and left to stir overnight at room temperature. The next day, sodium chloride was added to the reaction to obtain a 2 M final concentration. The solution was dialyzed (6 – 8K MW), alternating between DI water and 1 M NaCl solution and ending with DI water for at least 2 days. The solution was then filtered through a 0.2 μm membrane, after which it was lyophilized and stored at -20°C until use. Proton NMR shifts, detected using H’ NMR, of pendant norbornenes in D_2_O, δ6.33 and δ6.02 (vinyl protons, endo), and δ6.26 and δ6.23 ppm (vinyl protons, exo), were compared to the heparin methyl group δ2.00 ppm to determine functionalization. Heparin-norbornene was confirmed by ^1^H NMR with 45.1% norbornene modification. All equivalents are based on the moles of the heparin repeat unit.

### nH Nanoparticle Fabrication

Modified heparin was used to form nanoparticles as previously described.^[26b, 27]^ Briefly, 100 mg heparin-ABH was dissolved at 100 mg/mL in sodium acetate solution (pH 4), mixed with 123 μL Tween-80 and 383 μL Span-80 in 10 mL hexane, and sonicated to form nanoparticles via radical polymerization initiation by addition of 60 μL of 300 mg mL^−1^ ammonium persulfate (APS) and 10 μL tetramethylethylenediamine (TEMED) as the radical initiator and catalyst. For heparin-NB, 100 mg lyophilized hep-NB was dissolved in 500 μL 0.3 M 2-[4-(2-hydroxyethyl)piperazin-1-yl]ethanesulfonic acid (HEPES, pH 8.2) with 0.27 mg dithiothreitol (DTT), and 129 μL of 50 mM tris(2-carboxyethyl)phosphine (TCEP) added. Heparin-NB solution was then added similarly to 10 mL hexane with 123 μL Tween-80 and 383 μL Span-80 and then sonicated to form nanoparticles via inverse emulsion. Radical initiation was similarly done by addition of APS and TEMED. After sonication, reaction solutions were allowed to stir overnight and in the dark. ABH- and norbornene-modified heparin nanoparticles (nH-ABH or nH-NB) were purified with a liquid-liquid extraction. 40 mL of hexane, 10 mL of brine, and heparin nanoparticle mixture were first added to a separatory funnel. The bottom aqueous phase layer contained brine and heparin nanoparticles, which was extracted 7 times with 40 mL of hexane until clear. The aqueous nanoparticle solution was then dialyzed against against DI water for 2 days in 3500 MW dialysis tubing and then 0.2 μm sterile filtered before storing at 4 °C until use.

### Heparin Nanoparticle Characterization

Heparin nanoparticle (nH) diameter was determined by both dynamic light scattering (DLS) and scanning electron microscopy (SEM), as previously described.^[26b, 27]^ For DLS, nH were loaded into a clean cuvette which was then loaded into a Anton Paar Litesizer DLS machine for at least 10 runs, with each run comprised of three measurements. Data was reported as relative frequency intensity weighted particle diameter distribution, mean diameter, and polydispersity (PDI). For SEM, nH was serially diluted and added to dry on round glass coverslips prior to addition of a thin layer of gold via Denton Desk V sputter coater. Gold coated nH samples were then imaged in Hitachi TM3030Plus Tabletop SEM.

### VEGF + nH Binding (CLUVENA Fabrication)

Clustered VEGF nanoparticles (CLUVENA) were created by covalently tethering vascular endothelial growth factor (VEGF) to nH as previously described.^[26b, 27]^ Briefly, human recombinant VEGF-165 (BioLegend, 583706) was added to nH-NB in 10 mM LAP in 1x PBS at 2 μg VEGF per 0.1 μg nH in low binding protein tubes. VEGF and nH-NB solution was incubated on ice for 30 min before being illuminated with 20mW/cm^2^ of UV light (365 nm) for 30 min. CLUVENA was then transferred to a 1% bovine serum albumin (BSA) blocked 100 kDa molecular weight cut-off Amicon centrifugal filter to remove unbound VEGF at 14,000 RPM for 10 min. After each spin, 500 μL of 0.05% Tween-20 in PBS was added. This process was repeated four times. Next, 500 μL of 1x PBS was added and then spun down at 14,000 RPM for 10 min. This process was repeated three times. On the last spin, the Amicon filter was flipped over and spun down at 14,000 RPM for 15 min to collect CLUVENA. The initial flow through and the subsequent wash flow through solutions were saved to calculate the final VEGF concentration remaining bound to CLUVENA.

### VEGF ELISA

The binding efficacy of VEGF to nanoparticles was measured using a standard VEGF ELISA of flowthrough and washes as previously described.^[27, 30]^ VEGF ELISA (R&D Systems, DY293B-05) was performed following the manufacturer’s instructions. A VEGF standard curve from 2 ng mL^−1^ to 0 ng mL^−1^ was sufficient while initial flowthrough and first wash were diluted 1000x, subsequent 0.05% Tween-20 PBS washes were diluted 100x, and PBS washes were diluted 10x to be within limit of detection.

### VEGFR-2 Phosphorylation Assay

Human Umbilical Vein Endothelial Cells (HUVECs, Duke University, Cell Culture Facility) were cultured in EGM-2 media (Lonza) and tested for VEGFR-2 phosphorylation as previously described.^[53]^ Briefly, HUVECs were starved with serum-free media for 6 hours. Cells were then treated with CLUVENA and controls in sterile 1x PBS for 15 minutes. After treatment, cells were washed, lysed and sufficiently diluted to run Total VEGFR-2 and phospho-VEGFR-2 ELISA (R&D Systems, Minneapolis, MN). Ratio of phosphorylated-to-total VEGFR-2 indicates degree of VEGFR-2 activation.

### Modification of Hyaluronic Acid with Norbornene

Hyaluronic acid-norbornene (HA-NB) was synthesized as previously described.^[26b]^ 80 mL of 200 mM MES buffer, prepared at a pH of 5.5, was used to dissolve 1 g of 79,000 Da MW HA. Once the HA was completely dissolved, 3.1 g (4 molar equivalents) of 4-(4,6-Dimethoxy[1.3.5]triazin-2-yl)-4-methylmorpholinium chloride (DMTMM) (MW: 294.74 Da) (TCI America, Portland, OR) was added. The reaction was then allowed to stir for 10 min, after which 0.677 mL (2 molar equivalents) of 5-Norbornene-2-methanamine (NMA) (TCI America, Portland, OR) was added dropwise into the mixture. The reaction was left to stir at room temperature overnight. The next day, the reaction mixture was precipitated in 1 L of cold EtOH (200 proof). The precipitates were then dialyzed against DI water for 30 minutes in 2 M brine solution followed by an additional 30 minutes in 1 M brine solution. This process was repeated 3 times, after which the precipitates were dialyzed for 24 hours against DI water. The final product solution was collected, lyophilized, and stored at -20°C until use. HA-NB had a 33.5% NB functionalization, as confirmed by ^1^H-NMR. Functionalization was determined by considering ^1^H-NMR shifts of pendant norbornenes in D_2_O, δ6.33 and δ6.02 (vinyl protons, endo), and δ6.26 and δ6.23 ppm (vinyl protons, exo) compared to the HA methyl group δ2.05 ppm. All equivalents were based on the moles of the HA repeat unit.

### Synthesis of Tetrazine Crosslinker

Tetra-polyethylene glycol-tetrazine (4arm-PEG-Tet) was synthesized as previously described.^[26b, 54]^ First, 0.5 mL of CDCl_3_ was used to dissolve 100 mg of 20,000 Da MW tetra-PEG-SH (NOF America, White Plains, NY) and 15 mg of methyltetrazine- PEG4-maleimide (MW: 514.53 Da) (Kerafast, Boston, MA) (maleimide/SH ratio of 1.05). 1 µL (0.5 molar equivalent) of triethylamine (TEA) was then added to the solution, which was then left to stir for 4 h at room temperature. The product was precipitated in 50 mL cold diethyl ether and confirmed by ^1^H-NMR.

### Synthesis of AlexaFluor Tetrazine

Alexa Fluor 488 C2-tetrazine (Alexa488-Tet) was synthesized as previously described^[26b, 54]^. 0.17 mL CDCl_3_ was used to dissolve 2.8 mg (1 molar equivalent to Alexa Fluor C2 maleimide) of 3500 Da MW HS-PEG-SH (JenKem Technology USA, Plano, TX) and 0.41 mg (1 molar equivalent to Alexa Fluor C2 maleimide) of methyltetrazine-PEG4- maleimide (MW: 514.53 Da) (Kerafast, Boston, MA) . 0.11 µL (0.5 molar equivalent) of triethylamine (TEA) was then added to the mixture, after which the reaction was left to stir overnight at room temperature. The next day, 0.17 mL of CDCl_3_ was used to dissolve 1 mg of Alexa Fluor 488 C2 maleimide (MW: 1250 Da) (Thermo Fisher Scientific), which was then added to the reaction. 0.11 µL (0.5 molar equivalent) of triethylamine (TEA) was then added. The final reaction was left to stir overnight at room temperature. The product was precipitated in 10 mL of cold diethyl ether, dried under vacuum overnight, dissolved in dimethylformamide at 1 mg/mL, and then stored at −20 °C until use.

### Microgel Production and Purification

HA-NB microgels were produced as described previously.^[34]^ First, 1 mL of HA-NB gel precursor solution was prepared by dissolving lyophilized HA-NB product (final concentration = 3.5% w/v) in 50 mM HEPES (pH 7.5), after which di-thiol MMP sensitive linker peptide (Ac-GCRDGPQGIWGQDRCG-NH2, Genscript) (SH/HA ratio of 14), tris(2-carboxyethyl)phosphine (TCEP) (Sigma-Aldrich) (TCEP/SH ratio of 0.25), and lithium phenyl(2,4,6-trimethylbenzoyl)phosphinate photo-initiator (LAP) (final concentration = 9.90 mM) (TCI America, Portland, OR) were then added. The final solution was first filtered through a 0.22 µm sterile filter after which it was then transferred to a 1 mL BD Leur-Lok syringe. A 5 mL BD Leur-Lok syringe was filled with a 5% (v/v) Span-80 solution prepared in heavy mineral oil. Using tubing, the 1 mL syringe of the gel precursor solution and the 5 mL syringe of the 5% Span-80 solution were attached to the inner and outer inlets of a planar flow-focusing microfluidic device, respectively. A single syringe pump was used to simultaneously push both syringes at asymmetric flow rates at a flow rate of ∼6.4:1 (oil:aqueous), as previously described.^[62]^ To form microgels, a flow focusing region within the microfluidic device allowed the precursor solution to be “pinched” by the oil and the resulting droplets were crosslinked by exposure to UV light (20 mW/cm2) off-chip using an OmniCure LX500 LED Spot UV curing system controller with a OmnniCure LX500 LED MAX head (365nm wavelength, 30% power). A 15 mL conical tube wrapped in foil was used to collect the hydrogel microparticle (microgel) emulsion.

The microgel suspension was centrifuged at 5250 x g for 5 min, after which the material was maintained under sterile conditions. In a sterile hood, the supernatant oil was aspirated off and the remaining microgels were washed with sterile filtered HEPES buffer and centrifuged again at 5250 x g for 5 min. Washing was repeated until no more visible oil remained in the supernatant. Prior to use, particle endotoxin levels were determined with the Pierce LAL Chromogenic Endotoxin Quantitation Kit (Thermo Fisher Scientific) following the manufacturer’s instructions and were found to be consistently below 0.2 endotoxin U/mL. Particles were stored at 4°C until use.

### Microgel Post-Fabrication Modification

HA-NB microgels were modified post-fabrication with AF 488 and RGD-cell adhesion peptide as described previously^[34]^. 0.005 mM of Tet-AF 488 in 0.3 M HEPES buffer and 1 mM of thiolated RGD-cell adhesion peptide (RGDSPGERCG; Genscript, Piscataway, NJ) with 10 mM LAP in HEPES buffer was reacted with excess norbornene groups on the microgels for at least 1 hour. The suspended functionalized microgels were then pelleted by centrifuging at 14,000 x g for 5 min, after which they were washed three times with 0.3 M HEPES and recovered with the same centrifugation conditions.

### Generation of MAP Scaffolds and Characterization

An inverse-electron-demand Diels-Alder tetrazine-norbornene click reaction was used to crosslink between individual microgels, creating MAP scaffolds. In brief, excess norbornene groups on microgels were linked by 4arm-PEG-Tet to form tetrazine mediated MAP (Tet-MAP) scaffolds at an annealing ratio (HA monomer/tetrazine) of 7.

### Rheology

50 μL of freshly prepared MAP solution was sandwiched between 2 sigma-coated (Sigma Aldrich, St. Louis, MO) slides with 1 mm Teflon spacers, fastened with binder clips, and incubated for 2 hr at 37 °C to generate MAP scaffold discs 8 mm in diameter and 1 mm in height for rheological testing. All rheological measurements were performed with a plate-to-plate rheometer (Physica MCR, Anton Paar, Ashland, VA). A frequency sweep was performed using a strain of 1% and angular frequency ranging from 0.1 to 10 s^-1^.

### Void Volume Fraction (VVF)

Microgels labeled with AF-488 were used to make scaffolds similar to rheology preparation. MAP scaffolds were then incubated with TRITC-dextran 70kDa and imaged on Nikon C2 confocal microscope at z-step of 5 μm across 100 – 300 μm. Void volume fractions (VVF) were calculated by measuring volume of surface renderings on IMARIS (Oxford Instruments, Morrisville, NC) for AF488 labelled microgels and TRITC-dextran labeled void space. VVF from microgels subtracted microgel area divided by total scaffold volume from 1 whereas VVF from FITC-dextran used FITC-dextran volume divided by total scaffold volume.

### LOVAMAP Analysis

To characterize the non-particle void space of our MAP scaffolds, we use LOVAMAP, an in-house software designed to analyze granular material.^[36]^ For microscope image samples, LOVAMAP requires a pre-processing step that converts 2D z-stacks into 3D data and identifies the location of each particle. To accomplish this, we used custom software for 3D particle segmentation, which relies on interpolation and watershed to reformat data and segment particles.^[55]^ Processed data was then fed into LOVAMAP, and 3D pores are identified, which represent open 3D spaces between particles. Briefly, LOVAMAP accomplished this by using the location of particles as a basis for identifying medial axis landmarks, which were then used to delineate the open spaces separated from one another by narrow spaces. Since our microscope image samples are thin, we did not implement the feature of LOVAMAP that considers open spaces at the entrance / exit locations of the sample. Therefore, pore segmentation extends from the inside of the scaffold to the surface, which may lead to over-segmentation at the surface. We reported data for all 3D pores – both interior and those that lie at the surface of the sample. To characterize the void space of our materials, we reported distributions for 3D pore volume (pL), aspect ratio, and diameter of the largest enclosed sphere (µm).

### VEGF Release Assay

200 ng of VEGF (bound to heparin nanoparticles), 50 μL HA-NB microgels, and 4-arm PEG-tetrazine crosslinker (at Tetrazine/HA ratio of 7) were mixed to create a MAP-CLUVENA scaffold. The scaffold was incubated in 1x PBS at 37 °C for 11 days, during which PBS surrounding the scaffold was sampled at multiple timepoints. VEGF released into the surrounding 1x PBS was measured using ELISA (R&D Systems, DY293B-05) following manufacturer’s instructions.

### Photothrombotic Stroke

Animal procedures were performed in accordance with the US National Institutes of Health Animal Protection Guidelines and approved by the Chancellor’s Animal Research Committee as well as the Duke Office of Environment Health and Safety. C57BL/6 male mice of 8–12 weeks (Jackson Laboratories), were used in the study. A photothrombotic (PT) was performed as previously described.^[26b]^ Animals were put under isoflurane anesthesia (3% induction and 1.5% maintenance at 1 L min^−1^ USP grade O2). and placed on stereotactic apparatus. After application of eye ointment and sterilization of mice head, a midline incision was done to expose skull and connective tissue were removed. Laser was positioned 1.5 mm lateral from the bregma, and mouse was interperitoneally (IP) injected with photosensitive dye rose Bengal (Sigma; 10 mg mL^−1^ in PBS) at 10 μL g^−1^ mouse bodyweight. After 7 min, the brain was illuminated at 42 mW through intact skull for 13 min. A burr hole was drilled through the skull at 1.5 mm medial/lateral, same as laser position. The incision was then closed using Vetbond (3M, St. Paul, MN). Mice were placed in a clean cage on heat pad and monitored until awake and alert.

### Stereotactic Injection

5 days post-stroke, mice were treated with either 1x PBS, MAP + CLUVENA + nH, or MAP + CLUVENA. In brief, mice were first anesthetized with isoflurane at 3% and then transferred and positioned on a stereotactic stage for isoflurane maintenance at 1.5%. Following alcohol and iodine washes (3x), the skin covering the skull was re-opened, exposing the location of the photo-thrombotic stroke. 6 μL of treatment was injected directly into the stroke cavity through a flat 30-gauge needle attached to a 25 μL Hamilton syringe (Hamilton, Reno, NV). A syringe pump was used to deliver treatments at an infusion rate of 0.6 μL min^−1^ at the same coordinates as the PT stroke laser irradiation, 1.5 mm lateral from bregma, and 0.75 mm ventral from the skull. Prior to injection, the syringe was lowered 0.75 mm into the infarct site. 5 min after completion of the injection, the syringe was lifted out of the infarct site. The incision was then closed using Vetbond (3M, St. Paul, MN). Mice were placed in a clean cage on heat pad and monitored until awake and alert.

### Anti-Xa (Heparin) Assay

8-16 week old C57BL/6 male mice were anesthetized with isoflurane, after which 50 μL of PBS, heparin, heparin-NB, nH, or CLUVENA was administered via a retro-orbital injection. After 5 min, blood was collected from the inferior vena cava in a syringe pre-loaded with 50 μL 3.2% sodium citrate as an anticoagulant. Additional sodium citrate was added to collected blood to achieve a final 1:9 sodium citrate to blood volume ratio. Whole blood was spun down for 10 minutes at 5,000 x g at room temperature. Pseudo-plasma was collected and spun down again under the same conditions. Plasma was collected and stored at -80 °C. Anti-Xa/heparin assay was performed using HemosIL Liquid Heparin assay on ACL TOP Analyzer per manufacturer’s instructions (Beckman Coulter, Brea, CA).

### Tissue processing

At terminal endpoints, mice were anesthetized with isoflurane, after which 50 μL of biotinylated Lycopersicon esculentum (tomato) Lectin (Vector Labs, Burlingame, CA) was administered via a retro-orbital injection. After 10 minutes, the mice were first perfused with at least 10 mL of ice cold 1x PBS followed by ice cold 4% (wt/vol) paraformaldehyde (PFA). Following perfusion, brains were extracted and placed in a 4% PFA solution for 2-12 hours at 4°C. Brains were then washed several times with 1X PBS, after which they were placed into a 30% sucrose solution prepared in 1X PBS for 3 days at 4 °C. Brains were cryosectioned into 30 μm slices, collected onto gelatin coated glass slides, and preserved at -80°C until staining.

### Immunostaining

In preparation for immunofluorescent (IF) staining, slides were warmed to room temperature. Slides were then washed in 1x TBS (Sigma, D8537) for 5 min three times. A blocking buffer, comprised of 1x TBS, 0.3% Triton-X, and 10% normal donkey serum (NDS), was used to block tissue samples at room temperature for 1-2 hours. Primary antibodies, prepared in blocking buffer, were then left to incubate with tissue overnight at 4°C. The next day, slides were washed in 1x TBS + 0.3% Triton-X for 10 mins three times. Secondary antibodies and DAPI (1:1000), prepared in blocking buffer, were then left to incubate with tissue for 2 hours at room temperature, protected from light. Slides were then washed quickly with 1x TBS, followed by three additional 10 min 1x TBS washes. Tissue was left to dry at room temperature for approximately 1-2 hours. To dehydrate tissue sections, slides were placed in a series of ethanol baths: 50%, 50%, 70%, 70%, 90%, 90%, 95%, 95%, 100%, and 100%. Slides were left in each bath for approximately 1 min, except for the final 100% ethanol bath, during which slides were left to incubate for at least 3 minutes. Following dehydration, tissue was then defatted in two xylene baths: 1 min. in the first bath and at least 5 min in the second bath. Slides were then mounted in DPX and left to dry overnight at room temperature. *Primary Antibodies:* 1:400 GFAP (Invitrogen, 13-0400), 1:250 NF200 (Invitrogen, N4142), 1:250 Iba1 (Wako Chemicals, 011-27991), 1:250 Iba1 (Wako Chemicals, 019-19741), 1:250 CD31 (Santa Cruz Biotech, sc-18916), 1:400 Tmem119 (Synaptic Systems, 400 004), 1:200 AQP4 (Sigma-Aldrich, A5971). *Secondary Antibodies:* 1:500 Alexa Fluor 555 (Rb, Invitrogen, A-31572), 1:500 Alexa Fluor 488 (Rt, Invitrogen, A-21208), 1:500 Alexa Fluor 555 (Rt, Invitrogen, A48270), 1:500 Alexa Fluor 647 (Rt, Invitrogen, A48272), 1:500 Alexa Fluor 488 (Gt, Abcam, ab150129), 1:500 Alexa Fluor 555 (Gt, Abcam, ab150130), 1:500 Alexa Fluor 555 (Guinea Pig, Invitrogen, A-21450), 1:500 Alexa Fluor 647 (Biotin, Invitrogen, S32357), 1:500 Alexa Fluor 555 (Biotin, Invitrogen, S32355), 1:500 Alexa Fluor 488 (Biotin, Invitrogen, S11223).

### Imaging and Image Analysis

All fluorescent images were taken with a Nikon Ti Eclipse scanning confocal microscope equipped with a C2 laser with 4x or 20x air objectives. Fiji/ImageJ was used to analyze fluorescent images. Images are represented as maximum intensity projections (MIP). FIJI/ImageJ was used to analyze fluorescent images. For positive area measurements, images were despeckled, thresholded, and measured at appropriate ROI. Normalization was done by dividing signal positive area by void space, obtained by thresholding gaussian blurred DAPI image at sigma 8. For distance measurements such as scar thickness or infiltration, at least 10 lines were averaged per imaged section, 2 sections analyzed per mouse. Vessel maturation analysis with AQP4+ in TL+ was measured by using TL positive area as an ROI mask applied in the AQP4 channel to obtain AQP4 positive area only within TL positive area.

### Behavioral Testing

*3-*Dimensional aligned neural network for computational animal ethology (DANNCE) set up, software, hardware, and recordings were done as previously described for 6 cameras.^[50]^ Briefly, cameras were calibrated to intrinsically and extrinsically. Camera intrinsics were computed and calibrated using AprilTags.^[56]^ Intrinsic calibration was repeated if mean pixel error is greater than 0.5 pixel or each camera has less than 150 frames with detectable AprilTags. Camera extrinsics were calibrated by labeling four points on objects placed in the recording environment of known positions and dimensions. Animal motion capture recordings were done by placing mice into 4” diameter glass cylinder and recorded for 5 min at 100 fps. In between mice, stage and cylinders were wiped down with 70% ethanol. Animal recordings were consistently done in the afternoon to evening. Video predictions were done by DANNCE algorithm as described previously.^[50]^ DANNCE generates 3-D outputs for 20 skeletal keypoints based on real-world coordinates. Keypoints of the forepaws are used to analyze the number of frames corresponding to rearing events. Given the known coordinates of the cylinder’s center, the distance between the cylinder and the forepaws is calculated. Frames are classified as rearing events if the forepaws are positioned at least 30 mm above the ground and the distance between the forepaws and the cylinder is less than 15 mm. The final outcome is expressed as the proportion of frames identified as rearing events relative to the total number of frames in each video, normalized by the baseline measurement.

### Statistical Analysis

For immunofluorescent image quantification, data is presented in floating bar (min to max) where each point represents a single animal whereby values from two sections were averaged. Image quantifications are also presented as aligned dot plot representing the group mean where error bars represent standard deviation. One-way ANOVA was performed on quantification images which yielded P < 0.05, prompting a post-hoc analysis (Tukey HSD). To compare across time within treatment groups, two-way ANOVA was performed on images which yielded an interaction term P < 0.05, prompting a post-hoc analysis (Sidak test). Animals were excluded from analyses only when tissue was too damaged to analyze. For behavioral assessment, data is presented with error bars representing standard error where each point represents a single animal as biological replicate. Mixed-effect analysis was performed on cylinder test data which yielded P < 0.05 for both time and treatment factors. A Sidak post-hoc test was then performed comparing treatments group to stroke group within each timepoint. Animals were excluded from analyses only when they died or were sacrificed early due to humane endpoint. For all experiments, significance is indicated by *P < 0.05, **P < 0.01, ***P < 0.001, ****P < 0.0001. Statistical analyses were performed with GraphPad Prism software.

## Supporting Information

Supporting Information is available.

## Acknowledgements

The authors would like to acknowledge the following funding sources: the National Institutes of Health and the National Institute of Neurological Disorders and Stroke (R01NS079691), the National Institutes of Health (T.W.D.), and the National Institute on Drug Abuse (R34DA059512) (T.W.D.). N.I.J. was supported by an NIH Research Supplement to Promote Diversity in Health-Related Research (NS079691). This work was performed in part at the Duke University Shared Materials Instrumentation Facility (SMIF), a member of the North Carolina Research Triangle Nanotechnology Network (RTNN), which is supported by the National Science Foundation (award number ECCS-2025064) as part of the National Nanotechnology Coordinated Infrastructure (NNCI). The authors would like to thank Maria Notini from Duke Central Automated Laboratory (DCAL) for their expertise and assistance in coagulation testing.

## Notes

### Competing Interest Statement

TS is a founder of Tempo therapeutics which aims to commercialize MAP technology

