## Supplemental Information for "Clustered VEGF Nanoparticles in Microporous Annealed Particle (MAP) Hydrogel Accelerates Functional Recovery and Brain Tissue Repair after Stroke"

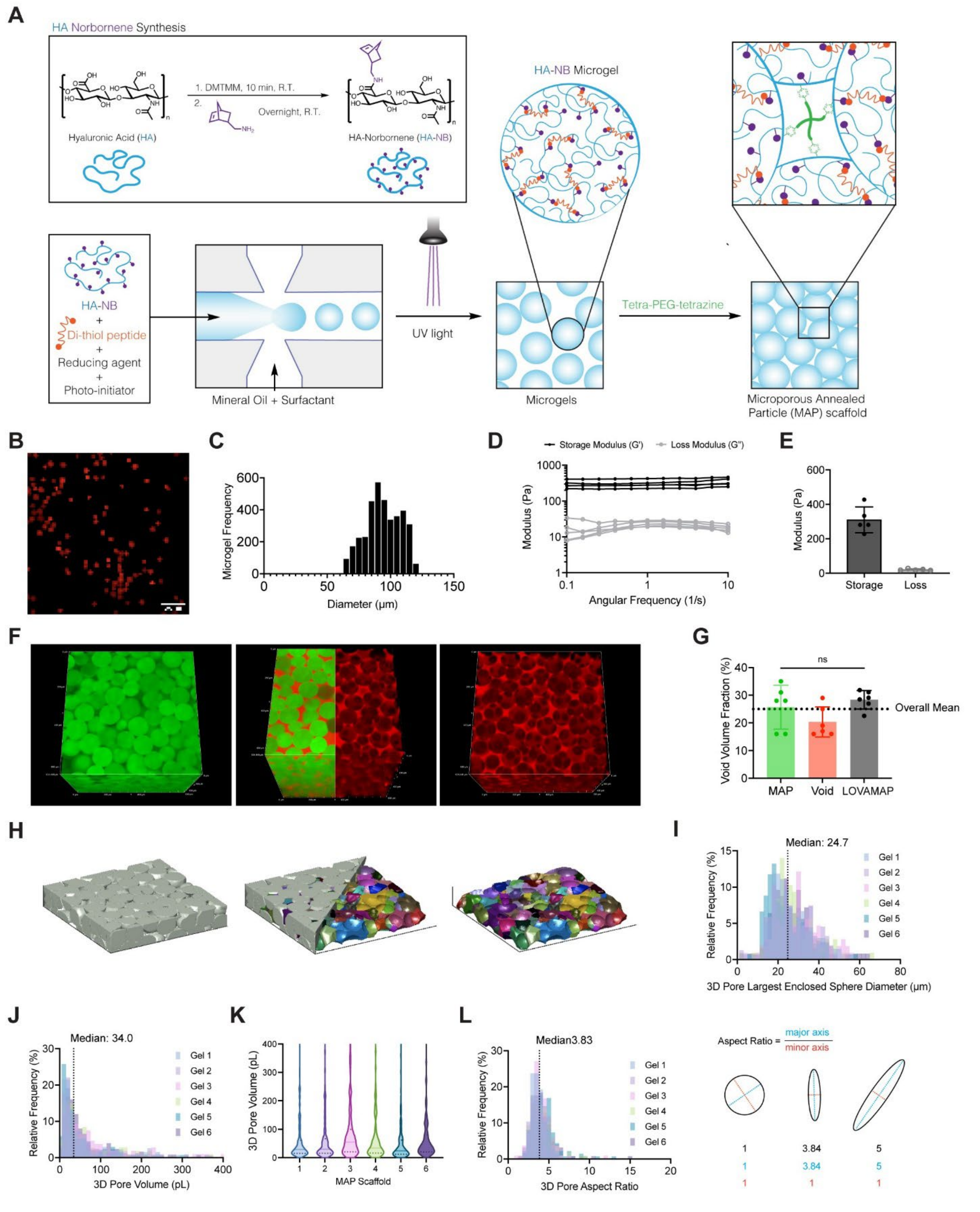


**Supplemental Figure 1.** HA-norbornene microgel and scaffold fabrication and characterization. A) Schematic of HA-norbornene (HA-NB) synthesis, microgel fabrication, and MAP formation. B) Representative fluorescent image of HA-NB microgels. C) Quantification of HA-NB microgel diameter, measured from fluorescent images of microgels for total of 3613 microgels. D)

Frequency sweep and E) storage and loss modulus of MAP scaffold at crosslinking ratio (Tetrazine/HA) of 7. F) Nikon volume rendering of MAP scaffold (green) and void fraction (red). G) Quantification of void volume fraction (%) from IMARIS renderings. H) Rendering of MAP scaffold highlighting the interior 3D pore space. All distinct colors represent a unique 3D pore. LOVAMAP quantification of I) the relative 3D-pore largest enclosed sphere diameter, J-K) 3D pore volume, and L) the 3D pore aspect ratio. For panel G, one-way ANOVA with a Tukey HSD post-hoc analysis was performed. * $p < 0.05$, ** $p < 0.01$, *** $p < 0.001$, **** $p < 0.0001$.

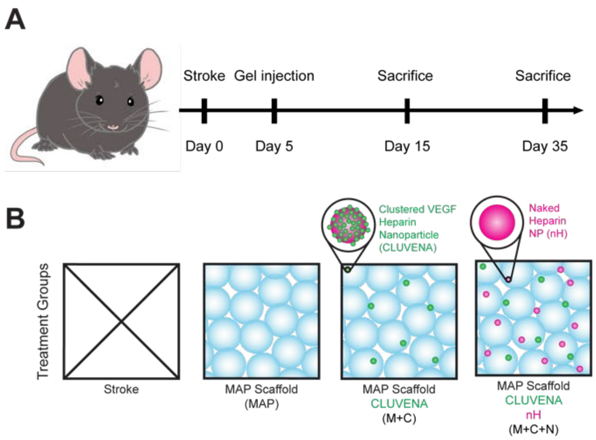


**Supplemental Figure 2.** Animal study design. A) Schematic of in vivo study timeline with PT stroke at day 0, gel injection at day 5, and sacrifice timepoints day 15 and day 35. B) Treatment groups including stroke-only control, MAP scaffold, MAP + CLUVENA (M+C), and MAP + CLUVENA + nH (M+C+N). C) Schematic of tissue processing and image analysis including an example of 20x tiled image with a representative image zooming into infarct. Tissue is stained for vessels (tomato lectin) in white, gel in green, and nuclei (DAPI) in blue.

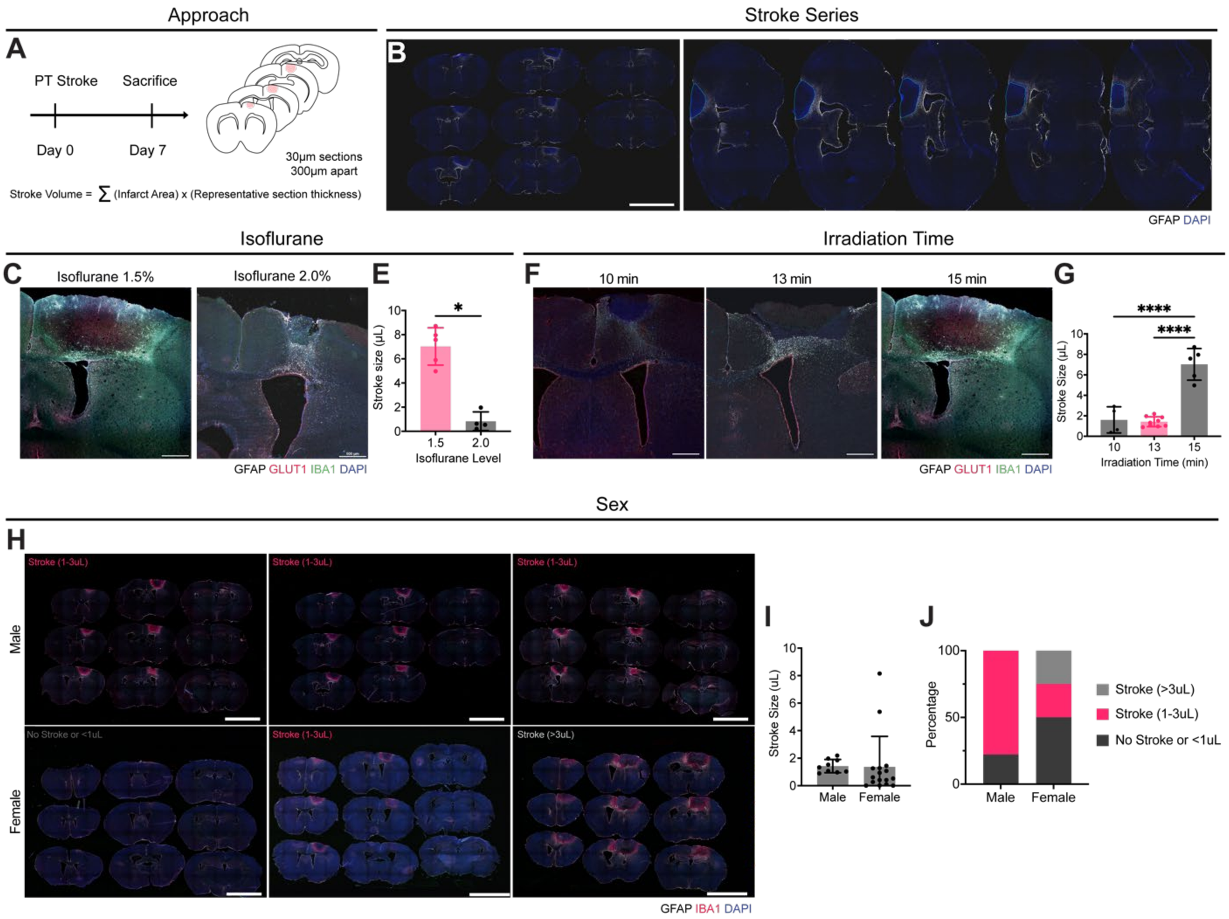


**Supplemental Figure 3.** Photothrombotic stroke baseline optimization. A) Experimental timeline and image analysis schematic. B) Representative immunofluorescent images of a coronal section series showing the progression of the infarct area through different regions of the brain. C) Representative immunofluorescent images of the infarct site depicting the effect of the delivery of different levels of isoflurane on the stroke size. Tissue is stained for astrocytes (GFAP) in white, glucose transporter 1 (GLUT1) in red, microglia (Iba1) in green, and nuclei (DAPI) in blue. D) Comparison of stroke volume, measured by a summation of glial scar barrier area measurements across coronal sections, between mice given a PT stroke while maintained under either 1.5% or 2.0% isoflurane. E) Representative immunofluorescent images of the infarct site depicting the effect of laser irradiation time during PT stroke on the final stroke size. Tissue is stained for astrocytes (GFAP) in white, glucose transporter 1 (GLUT1) in red, microglia (Iba1) in green, and nuclei (DAPI) in blue. F) Comparison of stroke volume, measured by a summation of glial scar barrier area measurements across coronal sections, between mice irradiated for 10, 13, or 15 mins. G) Representative immunofluorescent images of the infarct site in male and female mice. Tissue is stained for astrocytes (GFAP) in white, microglia (Iba1) in red, and nuclei (DAPI) in blue. H)

Comparison of stroke volume, measured by a summation of glial scar barrier area measurements across coronal sections, between male and female mice. I) For B,H, scale bar = 5000 µm. For panel D, a paired t-test was performed. For panel F, one-way ANOVA with a Tukey HSD post-hoc analysis was performed. For panel H, an unpaired t-test was performed. Error bars represent standard deviation. $n = 9 – 16$. * $p < 0.05$, ** $p < 0.01$, *** $p < 0.001$, **** $p < 0.0001$.
